# Spatially patterned excitatory neuron subtypes and circuits within the claustrum

**DOI:** 10.1101/2021.04.21.440755

**Authors:** Sarah R. Erwin, Brianna N. Bristow, Kaitlin E. Sullivan, Brian Marriott, Lihua Wang, Jody Clements, Andrew L. Lemire, Jesse Jackson, Mark S. Cembrowski

## Abstract

The claustrum is a functionally and structurally complex brain region, whose very spatial extent remains debated. Histochemical-based approaches typically treat the claustrum as a relatively narrow region that primarily projects to the neocortex, whereas circuit-based approaches suggest a broader region embedding neocortical and other neural circuits. Here, we took a bottom-up, cell-type-specific approach to complement and possibly unite these seemingly disparate conclusions. Using single-cell RNA-sequencing, we found that the claustrum is comprised of two excitatory neuron subtypes that are differentiable from the surrounding cortex. Multicolor retrograde tracing in conjunction with 12-channel multiplexed *in situ* hybridization revealed a core-shell spatial arrangement of these subtypes, as well as differential projection targets. Thus, the claustrum is comprised of excitatory neuron subtypes with distinct molecular and circuit properties, whose spatial patterns reflect the narrower and broader claustral extents debated in previous research. This subtype-specific heterogeneity likely shapes the functional complexity of the claustrum.

## INTRODUCTION

The claustrum has been implicated in a variety of functions and behaviours, including attention (Atlan et al., 2018; Goll et al., 2015; Smith et al., 2019), impulsivity (Liu et al., 2019), sleep (Narikiyo et al., 2020; Norimoto et al., 2020; Renouard et al., 2015), and the integration of information to support consciousness (Crick and Koch, 2005; Smythies et al., 2012). To determine the mechanistic contributions of the claustrum to these putative functions, it is essential to understand both the intrinsic organization of claustrum neurons, as well as how this organization relates to connectivity and function (Edelstein and Denaro, 2004). However, such an interpretation is challenged by the fact that even the precise anatomical boundaries of the claustrum are a matter of debate (Dillingham et al., 2019; Smith et al., 2019).

Here, utilizing a multimodal cell typing approach, we sought to understand the extent of heterogeneity within the excitatory claustrum neuron population, and relate this to local boundaries and long-range projections. Beginning with single-cell RNA sequencing, we identified two discrete populations of excitatory claustral neurons. To map the topography of these populations, we used multiplexed single molecule fluorescent *in situ* hybridization, revealing subtypes of core and shell claustral neurons that were transcriptomically distinct from surrounding cortical neurons. Combining this with multicolor retrograde tracing, we revealed a spatial organization of distinct cortical-projecting claustral populations that mapped onto the identified core and shell subtypes. This work demonstrates that the claustrum consists of heterogeneous populations of excitatory neurons that are topographically organized and project to functionally dissociable cortical regions, suggesting subtype-specific functionality of excitatory claustral neurons. To facilitate future research analyzing claustral cell-type-specific structure and function, data and analysis tools from this study are available via our interactive web portal (http://scrnaseq.janelia.org/claustrum; temporary username: reviewers; password: for_review).

## RESULTS

### scRNA-seq reveals discrete excitatory neuron subtypes within the claustrum

We began by using single-cell RNA sequencing (scRNA-seq) to understand the transcriptomic organization of the claustrum. From claustral microdissections from four mice, we manually harvested 1,112 cells based on a combination of unbiased blind selection of cells and selection of specific labeled projections (to either the lateral entorhinal cortex or retrosplenial cortex, “LEC” and “RSC” respectively; see Methods). After library preparation, sequencing, and filtering, we retained a total of 1,011 excitatory neurons for analysis (n = 478 cells blindly selected; n = 286 and 247 cells projecting to the LEC and RSC respectively).

We initially examined this dataset agnostic to any projection-specific information. Combining UMAP nonlinear dimensionality reduction (McInnes et al., 2018) with Louvain graph-based clustering (Stuart et al., 2019) revealed that cells broadly conformed to three transcriptomically separated clusters (Figure 1A; also seen in t-SNE: Figure 1 – figure supplement 1A). These clusters were all associated with expression of excitatory neuronal markers (Figure 1B), and were found across the anterior-posterior axis and across animals (Figure 1 – figure supplements 1B,C). In seeking to assign transcriptomic phenotypes to these cells, we noted one cluster (“Cluster 1”) was enriched for the claustrum marker gene *Synpr* (Binks et al., 2019; Wang et al., 2017). This cluster and a second cluster (“Cluster 2”) exhibited enriched expression of other claustrum marker genes (e.g., *Lxn, Gnb4*) relative to the third cluster (“Cluster 3”) (Figure 1C). Conversely, Cluster 3 was enriched for markers of excitatory cortical populations (e.g., the Layer 6b marker *Ctgf*, the Layer 6a marker *Sla*, and the general Layer 6 marker *Foxp2*) (Tasic et al., 2016; Tasic et al., 2018), suggesting a cortical phenotype (Figure 1C,D). Each cluster exhibited marker genes (Figure 1E; full lists: Supplemental Files 1-3), many of which were associated with neuronally relevant functions (Figure 1F; see Methods), suggesting higher-order structural and functional heterogeneity between these three clusters (full lists of differentially expressed genes: Supplemental Tables 1-3).

**Figure 1.**
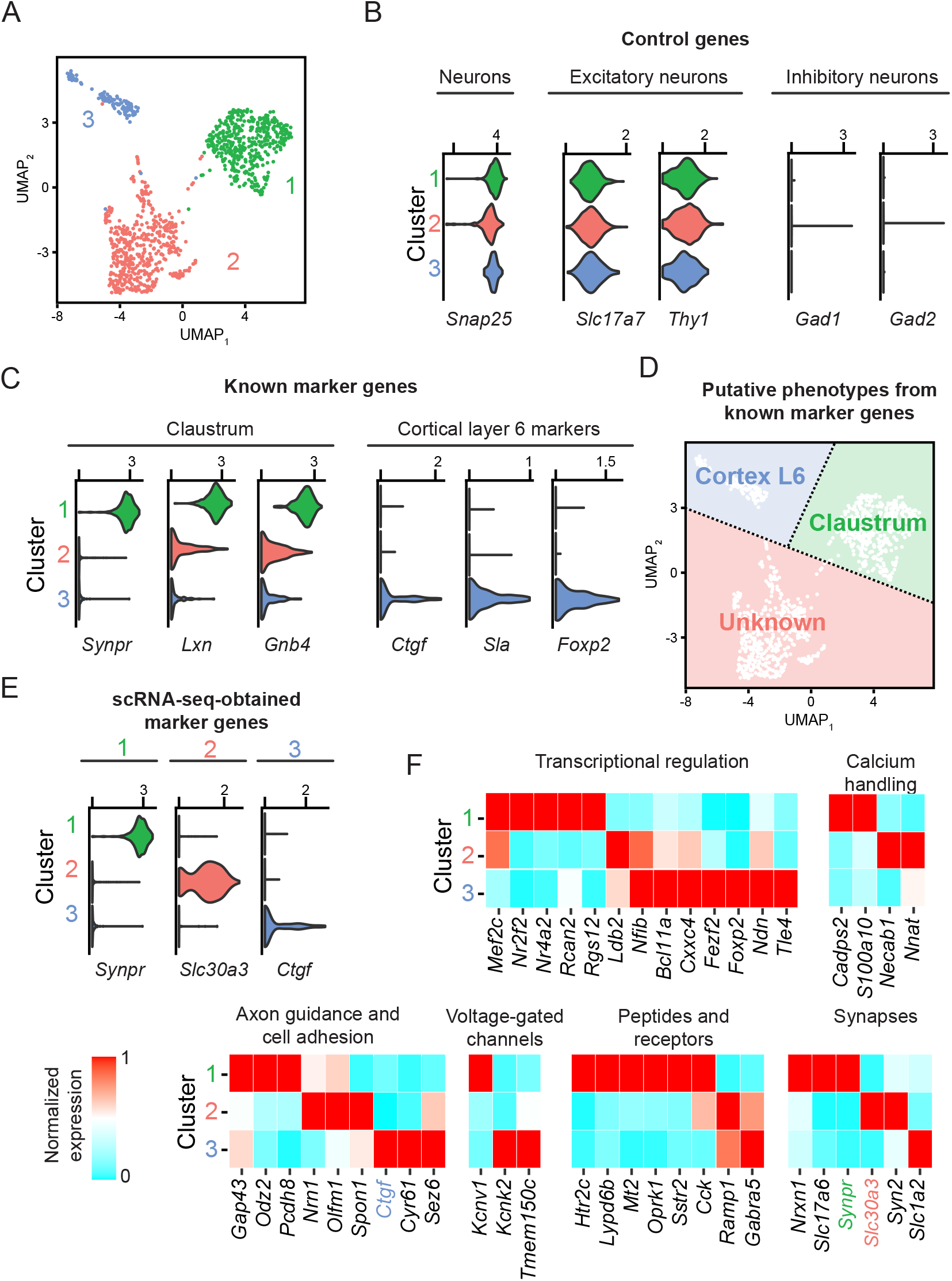
Excitatory claustrum-occupying cells are separable into discrete transcriptomic populations. A. UMAP dimensionality reduction of single-cell transcriptomes. Points denote individual cells, with coloring denoting cluster identity obtained by graph-based clustering. B. Violin plots illustrating expression of control marker genes, with accompanying values denoting normalized and log-transformed count value associated with right tick mark. C. As in (B), but for known marker genes of claustrum neurons and Layer 6 cortical neurons. D. UMAP graph annotated with putative subtypes. E. As in (B,C), but for cluster-specific marker genes derived from scRNA-seq data. F. Heatmap illustrating expression of genes associated with neuronal functionality that are enriched or depleted in a cell-type-specific fashion.

### Comparison to published work

To understand our results in the context of published scRNA-seq, we next compared our data to published datasets that potentially included the claustrum (Saunders et al., 2018; Zeisel et al., 2018). Consistent with the three clusters identified within our dataset, our dataset largely conformed to three distinct locations within the broader cell-type landscape when incorporating published data (Figure 1 - figure supplement 2A–B). In particular, Cluster 1 cells occupied an isolated group of *Synpr*-expressing cells, whereas Cluster 2 cells coarsely occupied a distinct location nearby other datasets, but were also enriched for specific marker genes like *Nnat* (Figure 1 - figure supplement 2C-J). In agreement with Cluster 2 being non-cortical, these *Nnat-* expressing Cluster 2 cells were also depleted in *Pcp4* expression (Figure 1 - figure supplement 3), a gene strongly expressed in deep cortical layers (Figure 1 - figure supplement 4) (but with exception of layer 6 intratelencephalic excitatory neurons: Watakabe et al., 2012). In collection, this work shows that our scRNA-seq data recapitulates previously described cell types, and further suggests new marker genes and cell types that may have gone underresolved in previous studies.

### Two types of excitatory claustral neurons exist in a core-shell arrangement

As the spatial cell-type-specific organization of the claustrum remains uncertain, we next sought to map our scRNA-seq-identified cell types in a spatial context. To do this, we used multiplexed fluorescent *in situ* hybridization (mFISH), allowing us to map 12 gene-expression targets at a single-molecule and single-cell resolution (O’Leary et al., 2020). We selected genes that allowed us to grossly identify excitatory neurons (*Slc17a7, Slc17a6*), cortical markers (*Ctgf, Pcp4*), classical claustrum markers (*Synpr, Lxn, Gnb4*), and new putative subtype-specific markers from our scRNA-seq dataset (*Cdh9, Slc30a3, Gfra1, Nnat, Spon1*) (overview of genes in scRNA-seq: Figure 2 - figure supplement 1).

We used mFISH to spatially register expression of these 12 genes across anterior, intermediate, and posterior sections of the claustrum (Figure 2A-D; expansions: Figure 2 - figure supplement 2; video: Supplementary File 4; n = 18,957 excitatory neurons from n = 5 animals analyzed). In doing so, we identified a claustrum population with a relatively central core-like location that exhibited expression of *Synpr*, and a surrounding shell-like population exhibiting expression of *Nnat*. This organization was present across the anterior-posterior axis (Figure 2 – figure supplement 3) and animals (Figure 2 – figure supplement 4), and recapitulated gene-expression properties expected from scRNA-seq (Figure 2 – figure supplement 5). Adjacent to these populations were other neuronal subtypes enriched for markers of cortical cells, including a cluster with spatial and transcriptional properties of deep layer 6 cells (i.e. a *Ctgf*-expressing cluster in the deepest cortical layer). Collectively, these results illustrated a core-shell organization of claustrum excitatory neuron subtypes surrounded by cortical cells (Figure 2E).

**Figure 2.**
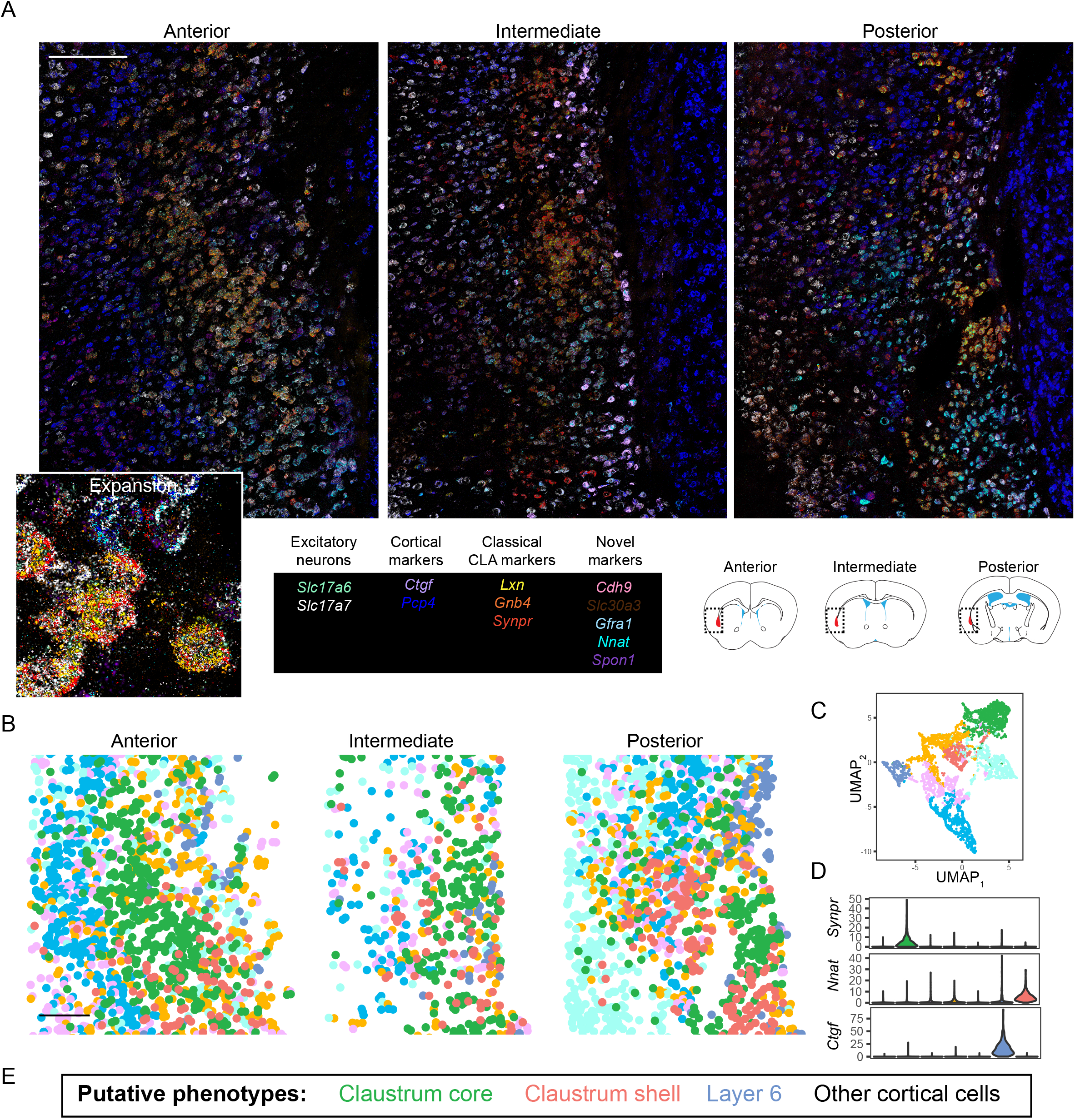
Multiplexed FISH analysis of the claustrum. A. Overview of anterior (left), intermediate (middle), and posterior (right) sections of the claustrum. Inset shows expansion of anterior section. Probe list provided at bottom middle, with atlas schematics denoting coronal section locations at bottom right, as well as imaged regions and claustrum definition of atlas (red). Scale bar: 200 μm. Atlas schematic adapted from (Franklin and Paxinos, 2013). B. *Slc17a7*-expressing cells (putative excitatory neurons) segmented from (A) and colored according to cluster identity for a representative animal. Scale bar: 200 μm. C. UMAP-based nonlinear dimensionality reduction for cells shown in (B). D. Expression of example marker genes for core clautrum (*Synpr*), shell clautrum (*Nnat*), and Layer 6 neurons (*Ctgf*). E. Putative phenotypes of cellular clusters, based upon marker gene expression in conjunction with spatial location.

### Claustrum excitatory subpopulations co-vary with projection target

Does this differential marker gene expression and spatial patterning correspond to distinct claustral circuits? To answer this question, we next considered projections to the retrosplenial cortex (RSC) and lateral entorhinal cortex (LEC), two claustral projections that exhibit minimal overlap (two-colour retrograde viral injections: Figure 3A; see also Marriott et al., 2020). We first examined our scRNA-seq dataset with respect to projection targets, where a subset of RSC- and LEC-projecting cells were specifically targeted by retrograde labeling and manual harvesting (Figure 3B). Strikingly, 84% (215/245) of RSC-projecting cells mapped onto the *Synpr*-expressing class, whereas 82% (229/281) of LEC-projecting cells mapped onto the *Nnat-*expressing class (Figure 3C). Similarly, applying mFISH to retrograde-labeled cells provided complementary evidence that *Synpr* and *Nnat* were respectively enriched in RSC-projecting and LEC-projecting cells (Figure 3D-I; other genes: Figure 3 – figure supplement 1), and illustrated that RSC- and LEC-projecting cells were enriched in distinct claustral subtypes (96% of RSC-projecting cells were found in core cluster and 54% of LEC-projecting cells were found in shell cluster, n = 4 and n = 2 animals respectively, Figure 3J). Thus, distinct excitatory claustrum projection neurons were coherently separable by marker genes, local spatial organization, and long-range projection targets.

**Figure 3.**
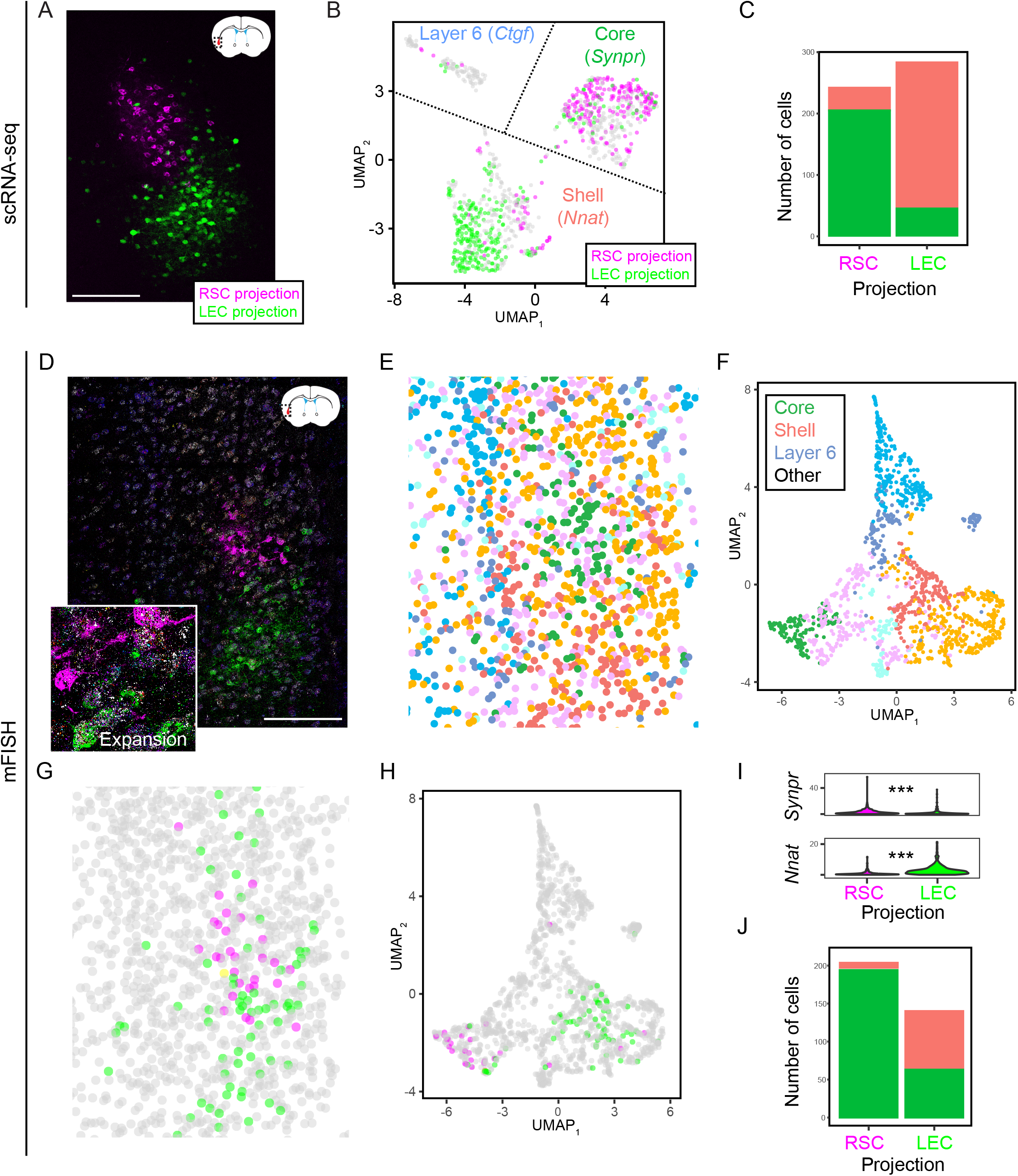
Claustrum transcriptomic subtypes are associated with different projections. A. Projections to the RSC (magenta) and LEC (green) emanate from different spatial locations. Atlas schematic denotes coronal section location, adapted from (Franklin and Paxinos, 2013). Scale bar: 200 μm. B. Left: UMAP visualization of scRNA-seq CLA transcriptomes, with coloring of individual cells corresponding to their associated projection. Labels denote cluster phenotypes and example marker genes. C. Counts of RSC-projecting and LEC-projecting cells according to scRNA-seq core and shell phenotypes. D. mFISH of intermediate section, including circuit mapping of long-range projections to the RSC (magenta) and LEC (green). Scale bar: 200 μm. E. Cellular segmentation and cluster identification based upon gene expression detected via mFISH. F. UMAP dimensionality reduction of mFISH profiles, coloured according to cluster identity. Putative phenotypes of clusters, based upon marker gene expression, are provided. G. Locations of neurons projecting to the RSC (magenta), LEC (green), or both (yellow). Scale bar: 200 μm. H. As in (G), but with projections shown in UMAP embedding. I. mFISH-derived expression of *Synpr* and *Nnat* in cells that project to either the RSC (magenta) or LEC (green). J. As in (C), but for mFISH core and shell phenotypes.

## DISCUSSION

A variety of different approaches have previously been used to established the anatomical definitions of the claustrum. Marker-based approaches using individual genes such as *Lxn*, *Gnb4*, and *Slc17a6* coarsely demarcate the boundaries of the adult mouse claustrum (Fodoulian et al., 2020; Kitanishi and Matsuo, 2017; Mathur et al., 2009; Peng et al., 2020; Wang et al., 2017; Watakabe et al., 2012), but it is unclear if these different genes all converge upon a monolithic cellular population or embody different claustrum cell types (and potentially include phenotypically cortical cells: Bruguier et al., 2020; Molnar et al., 2020; Puelles et al., 2016). Retrograde tracing from the cortex has been useful for identifying claustrum projection neurons (Marriott et al., 2020; Minciacchi et al., 1985; Watson et al., 2017; Zingg et al., 2018), but similarly it remains unknown whether circuit-labeled claustrum cells are intrinsically homogeneous.

Our approach here, integrating both genetic and circuit level approaches, identified two distinct claustrum cell types (Figure 1, 2) enriched for different long-range projection targets (Figure 3). It is important to note that RSC and LEC reflect the two most parallel claustrum projection classes (Marriott et al., 2020), and thus it is likely that other claustral projections comprise more mixed core-shell transcriptomic phenotypes. Ultimately, this suggests a claustral organizational scheme wherein discretely separate transcriptomic subtypes are biased – but not wholly separable – according to long-range projection target (Cembrowski and Menon, 2018).

Collectively, our results will allow subtype-specific claustral function to be assayed in future experiments by leveraging either marker genes or projection pathways (Cembrowski, 2019). Thus, our findings here will help to guide and inform observational and interventional experiments, and bridge claustrum cell-type identity, structure, and function. To facilitate such experiments and interpretations, we have hosted our scRNA-seq data online in conjunction with analysis and visualization tools (http://scrnaseq.janelia.org/claustrum; temporary username: reviewers; password: for_review). This web portal will help to identify how specific genes, cell types, and circuits mechanistically drive claustral function and behavior.

## METHODS

All procedures were approved by the University of British Columbia Animal Care Committee, the University of Alberta Health Science Laboratory Animal Services Animal Care and Use Committee, and the Janelia Institutional Animal Care and Use Committee.

### Retrograde tracer injections

Mature C57BL/6 mice of either sex were used for injections, and randomly assigned retrograde injection locations and tracers. Mice were administered carprofen via ad-libitum water 24 hours prior to surgery, and for 72 hours after surgery to achieve a dose of 5mg/kg. For surgery, mice were initially anesthetized using 4% isoflurane and maintained with 1.0-2.5% isoflurane. Mice were secured in a stereotaxic frame, with body temperature maintained through an electric heating pad set at 37°C, and lubricant was applied to eyes to prevent drying. Local anesthetic (bupivacaine) was applied locally under the scalp, and an incision along midline was made to access bregma and all injection sites. Craniotomies were marked and manually drilled using a 400μm dental drill bit according to stereotaxic coordinates. Pulled pipettes (10 - 20 μm in diameter) were back filled with mineral oil and loaded with virus or tracers. All injections were made using pressure injection, with 200 nL of retrograde tracer (Tervo et al., 2016) being injected. The skin was sutured after completing all injections and sealed. After allowing for sufficient time for retrograde labeling, mice were subsequently sacrificed for either histology, RNA sequencing, or mFISH processing, as described below.

### Single-cell RNA sequencing data acquisition and analysis

We used a manual capture approach to harvest cells from n = 4 mature male C57BL/6 mice. To facilitate microdissection of the claustrum, fluorescent tracers were used to delineate and grossly microdissect the claustrum from horizontal sections. In one animal, retrograde rAAV2-retro-GFP (Tervo et al., 2016) was injected into the anterior cingulate cortex to facilitate gross microdissection of the claustrum (Jackson et al., 2018), but not used to select for individual cells (i.e., GFP expression was used to microdissect the claustrum but cells were picked blind relative to GFP expression). In this animal, cells from separate anterior and posterior sections were obtained, allowing analysis of potential anterior vs. posterior differences in claustrum gene expression (Figure 1 – figure supplement 1B). To build the projection-specific dataset, for the remaining three mice, green and red retrobeads were respectively injected into the lateral entorhinal cortex and retrosplenial cortex, with this labeling used for gross microdissection as well as to select a subset of projection-specific cells for RNA-seq.

In all cases, manual purification (Hempel et al., 2007) was used to capture cells in capillary needles in approximately 0.1-0.5 mL ACSF cocktail, and subsequently placed into 8-well strips containing 3 mL of nuclease-free PBS-BSA (0.1% BSA). Each strip of cells was flash frozen on dry ice, then stored at −80°C until cDNA synthesis. Reverse transcription master mix (2 mL 5X buffer (Thermo Fisher Scientific), 2 mL 5M Betaine (Sigma-Aldrich, St. Louis, MO), 0.2 mL 50 mM E5V6NEXT template switch oligo (Integrated DNA Technologies, Coralville, IA), 0.1 mL 200 U/mL Maxima H- RT (Thermo Fisher Scientific), 0.1 mL 40 U/mL RNasin (Lucigen, Middleton, WI), and 0.6 mL nuclease-free water (Thermo Fisher Scientific) was added to the approximately 5.5 mL lysis reaction and incubated at 42°C for 1.5 hour, followed by 10 minutes at 75°C to inactivate reverse transcriptase. PCR was performed by adding 10 mL 2X HiFi PCR mix (Kapa Biosystems) and 0.5 ml 60 mM SINGV6 primer with the following conditions: 98°C for 3 minutes, 20 cycles of 98°C for 20 seconds, 64°C for 15 seconds, 72°C for 4 minutes, with a final extension step of 5 minutes at 72°C. Plates were pooled across the columns into an 8-well strip to yield approximately 250 mL pooled PCR reaction. From this, 100 mL of each well in the 8-well strip was purified with 60 mL Ampure XP beads (0.6x ratio; Beckman Coulter, Brea, CA), washed twice with 75% ethanol, and eluted in 20 mL nuclease-free water. Equal volume of each of the eight samples was pooled to create the plate-level cDNA pool for tagmentation, and the concentration was determined using Qubit High-Sensitivity DNA kit (Thermo Fisher Scientific).

Six hundred pg cDNA from each plate of cells was used in a modified Nextera XT (Illumina, San Diego, CA) library preparation, but using the P5NEXTPT5 primer and extending the tagmentation time to 15 minutes. The resulting libraries were purified according to the Nextera XT protocol (0.6x ratio) and quantified by qPCR using Kapa Library Quantification (Kapa Biosystems). Four plates were pooled together on a HiSeq 2500 Rapid flow cell reading 25 bases in read 1, 8 bases in the i7 index read, and 125 bases in read 2. Alignment and count-based quantification of single-cell data was performed by removing adapters, tagging transcript reads to barcodes and UMIs, and aligned to the mouse genome via STAR (Dobin et al., 2013).

In total, 1112 were manually harvested and underwent sequencing. Of these initial 1112 cells, 27 putative non-neuronal cells were excluded due to low expression of *Snap25* (CPM<0.001), and 74 additional cells were excluded due to low *Slc17a7* (CPM<1e-10). The remaining 1011 cells exhibit 5.2 ± 1.1 thousand expressed genes/cell from 142 ± 98 thousand reads/cell, mean ± SD. The relatively high abundance of excitatory neurons sampled owed both to the targeted approach for harvesting circuit-labeled cells, as well as the fact that excitatory neurons are relatively abundant relative to interneurons in the claustrum. No blinding or randomization was used for the construction or analysis of this dataset. No *a priori* sample size was determined for the number of animals or cells to use; note that previous methods have indicated that several hundred cells from a single animal is sufficient to resolve heterogeneity within excitatory neuronal cell types (Cembrowski et al., 2018a; Cembrowski et al., 2018b).

Computational analysis was performed in R (RRID:SCR_001905) (R Development Core Team, 2008) using a combination of Seurat v3 (RRID:SCR_007322) (Satija et al., 2015; Stuart et al., 2019) and custom scripts (Cembrowski et al., 2018a). To analyze our data, a Seurat object was created via *CreateSeuratObject*(min.cells = 3, min.features = 200), variable features identified via *FindVariableFeatures(selection.method=‘vst’,nfeatures=2000)* and scaled via *ScaleData()*. Data was processed via *RunPCA()*, *JackStraw(num.replicate=100), RunTSNE(), FindNeighbors(), FindClusters(),* and *RunUMAP(reduction=‘pca’),* with 30 dimensions used throughout the analysis. This processed Seurat object was then used for downstream analysis. Subpopulation-specific enriched genes obeying *p_ADJ_* < 0.05 were obtained with Seurat via *FindMarkers()*, where *p_ADJ_* is the adjusted *p*-value from Seurat based on Bonferroni correction. Functionally relevant differentially expressed genes were obtained using *FindMarkers()*, allowing for both cluster-specific enriched and depleted genes obeying *p_ADJ_* < 0.05, and manually identified for functional relevance. Raw and processed scRNA-seq datasets have been deposited in the National Center for Biotechnology Information (NCBI) Gene Expression Omnibus under GEO: GSE149495.

To integrate and compare our scRNA-seq data to previously published data, we downloaded data from two previous studies that broadly sampled cortical cells in the mouse brain (Saunders et al., 2018; Zeisel et al., 2018). From (Saunders et al., 2018), we downloaded frontal cortex data from F_GRCm38.81.P60Cortex_noRep5_FRONTALonly.raw.dge.txt.gz (from http://dropviz.org), and used a threshold of 16,000 transcripts/cell to extract 2,877 total cells. After screening against cells that lacked *Snap25* and/or *Slc17a7* expression, 2,842 putative excitatory neurons were retained for analysis (genes expressed/cell: 4.8 ± 0.6 thousand, mean ± SD; transcripts/cell: 16.5 ± 5.0 thousand, mean ± SD). We used a similar number of cells from (Zeisel et al., 2018), obtained from l6_r4_telencephalon_projecting_excitatory_neurons.loom (from http://mousebrain.org/loomfiles_level_L6.html): 3,151 cells were obtained using a threshold for 7500 transcripts/cell, with 3,141 cells retained after requiring *Snap25* and *Slc17a7* expression (genes expressed/cell: 3.7 ± 0.4 thousand, mean ± SD; transcripts/cell: 9.6 ± 2.0 thousand, mean ± SD). Integration of these published datasets with our dataset was done in Seurat v3 (Stuart et al., 2019) by creating a Seurat object incorporating all datasets, and then using *SplitObject()* to split according to original dataset, allowing each dataset to independently undergo normalization and variable feature selection (handled identically to our data). Integration anchors were subsequently identified (via *FindIntegrationAnchors()*) and used for integration (via *IntegrateData()*), using 30 dimensions. From here, integrated data underwent scaling, dimensionality reduction, and clustering identically to the method used for our data.

### Multiplexed fluorescent in situ hybridization data acquisition and analysis

Custom probes for multiplexed fluorescent *in situ* hybridization (mFISH) were purchased from Advanced Cell Diagnostics and were as follows: *Cdh9* (443221-T1), *Ctgf* (314541-T2), *Slc17a6* (319171-T3), *Lxn* (585801-T4), *Slc30a3* (496291-T5), *Gfra1* (431781-T6), *Spon1* (492671-T7), *Gnb4* (460951-T8), *Nnat* (432631-T9), *Synpr* (500961-T10), *Pcp4* (402311-T11), *Slc17a7* (416631-T12). mFISH was generally performed as previously implemented (O'Leary et al., 2020). Briefly, mature male mice were randomly selected for mFISH, and were deeply anesthetized with isoflurane and perfused with phosphate buffered saline (PBS) followed by 4% paraformaldehyde (PFA) in PBS. Brains were dissected and post-fixed in 4% PFA for 2-4 hr. Brain sections (20 μm) were made using a cryostat tissue slicer and mounted on glass slides. Slides were subsequently stored at −80°C until use. For use, the tissue underwent pretreatment and antigen retrieval per the User Manual for Fixed Frozen Tissue (Advanced Cell Diagnostics). All 12 probes with unique tails (T1-T12) were hybridized to the tissue, amplified, and the tissue counterstained with DAPI. Using cleavable fluorophores with unique tails (T1-T12), probes were visualized four at a time via an iterative process of imaging, decoverslipping, cleaving the fluorophores, and adding the next four fluorophores. Multiplexed FISH performed on tissue with viral tracing was first counterstained with DAPI, coverslipped with ProLongGold antifade mounting medium, then imaged. The tissue was decoverslipped by soaking in 4X SSC. Following this, standard multiplexed FISH protocol was followed, with the antigen retrieval step quenching all endogenous viral fluorescent protein signal and DAPI signal.

Multiplexed FISH images were acquired with a 63x objective on a SP8 Leica white light laser confocal microscope (Leica Microsystems, Concord, Ontario, Canada). Z-stacks were acquired with a step size of 0.45 μm for each imaging round. Final composite images are psuedocoloured maximum intensity projections, including brightness adjustments applied to individual channels uniformly across the entire image, with channels opaquely overlaying one another ordered from highest to lowest expression.

Processing of multiplexed FISH images generally followed our previously reported (O’Leary et al., 2020) FIJI (Schindelin et al., 2012) analysis pipeline. Briefly, the DAPI signal from each round was used to rigidly register probe signals across rounds, followed by nonlinear elastic registration via bUnwarpJ (Arganda-Carreras et al., 2010) to accommodate any nonlinear tissue warping due to decoverslipping. The individual nuclei were then segmented and dilated by a factor of 5 μm to include the surrounding cytosol. The signal from each probe was then binarized, by thresholding at the last 0.2-1% of the histogram tail, and then the number of pixels within ROIs selected from segmentation were summed and normalized to the pixel area of the cell and multiplied by 100. This in effect corresponded to percent area covered (PAC) of the optical space of a cell.

A total of 5 mature male C57BL/6 mice, each with an anterior, intermediate, and posterior section, were used for mFISH. Four mice had 200 nL retrograde viral injections into the RSC, two of which had additional retrograde viral injections into the LEC. The last remaining mouse had no viral injections. Across these five animals, 33,155 cells in total were imaged. To facilitate analysis of excitatory neurons specifically, a threshold of 1 PAC of *Slc17a7* was required for each cell to be included in analysis, resulting in 18,957 total putative excitatory neurons being used for subsequent analysis. *Slc17a7* expression levels were used only for cellular phenotyping, and thus excluded from further analysis.

For analysis, UMAP dimensionality reduction (McInnes et al., 2018) was performed on within-cell normalized CPA values using the *umap* package (15 nearest neighbors, all other parameters default), and hierarchical clustering performed with the *hclust* package using Euclidean distances between cells and Ward’s minimum variance method. In general, choosing 7 clusters was found to produce good agreement between UMAP dimensionality reduction and cluster assignments, although only 4 clusters were typically needed to separate cortical cell types from core and shell claustrum subtypes. For visualizations of representative individual animals across the anterior-posterior axis (Figure 2, Figure 2 – figure supplement 3) or individual sections in circuit mapping (Figure 3D-H), associated clustering and UMAP embedding shown reflect the illustrated tissue sections. For aggregate analysis (Figure 3I,J, Figure 2 – figure supplement 4), analysis includes all corresponding sections and animals. In general, violin plots show distribution of gene expression contingent on cluster identity or projection target, and Mann-Whitney U tests with a Bonferroni correction were used to identify differentially expressed genes. For correlating projection targets with mFISH results, cells labeled with fluorescent retrograde tracers were manually identified (n = 725 total across n = 4 animals), done in a blinded fashion relative to mFISH analysis. A small minority of cells that projected to both the LEC and RSC (1.9%: n = 14/725, consistent with Marriott et al., 2020) were excluded when comparing properties of LEC- vs. RSC-projecting neurons.

## ACKNOWLEDGEMENTS

MSC is supported by the University of British Columbia (Department of Cellular and Physiological Sciences, Djavad Mowafaghian Centre for Brain Health, and the Faculty of Medicine Research Office), the Natural Sciences and Engineering Research Council of Canada (RGPIN-2019-04507), the Canadian Institutes of Health Research (PJT-419798), and the Canadian Foundation for Innovation (John R. Evans Leaders Fund 38369). BM was supported by a studentship from the Neuroscience and Mental Health Institute. JJ is funded by the University of Alberta (Faculty of Medicine & Dentistry, and Department of Physiology), the Natural Sciences and Engineering Research Council of Canada (RGPIN-2018-05212), the Brain and Behavioral Research Foundation Young Investigator Grant, and Canadian Foundation for Innovation (John R. Evans Leaders Fund). This work was supported by resources made available through the NeuroImaging and NeuroComputation Centre at the Djavad Mowafaghian Centre for Brain Health (RRID: SCR_019086). Collaboration between MSC and LW, JC, and ALL was supported by the Janelia Visiting Scientist Program. We thank members of the Cembrowski lab for helpful discussions, and Jeffrey LeDue for insight and guidance in image acquisition.

## AUTHOR CONTRIBUTIONS

MSC and JJ designed and supervised the project, LW and AL collected single-cell RNA-seq data, SRE and BM performed retrograde tracing, JC constructed the web portal, SRE and BNB performed *in situ* hybridization, SRE, BNB, BM, JJ, and MSC analyzed the data, SRE, BNB, JJ, and MSC wrote the paper.

## SUPPLEMENTAL FIGURES

**Figure 1 - figure supplement 1.**
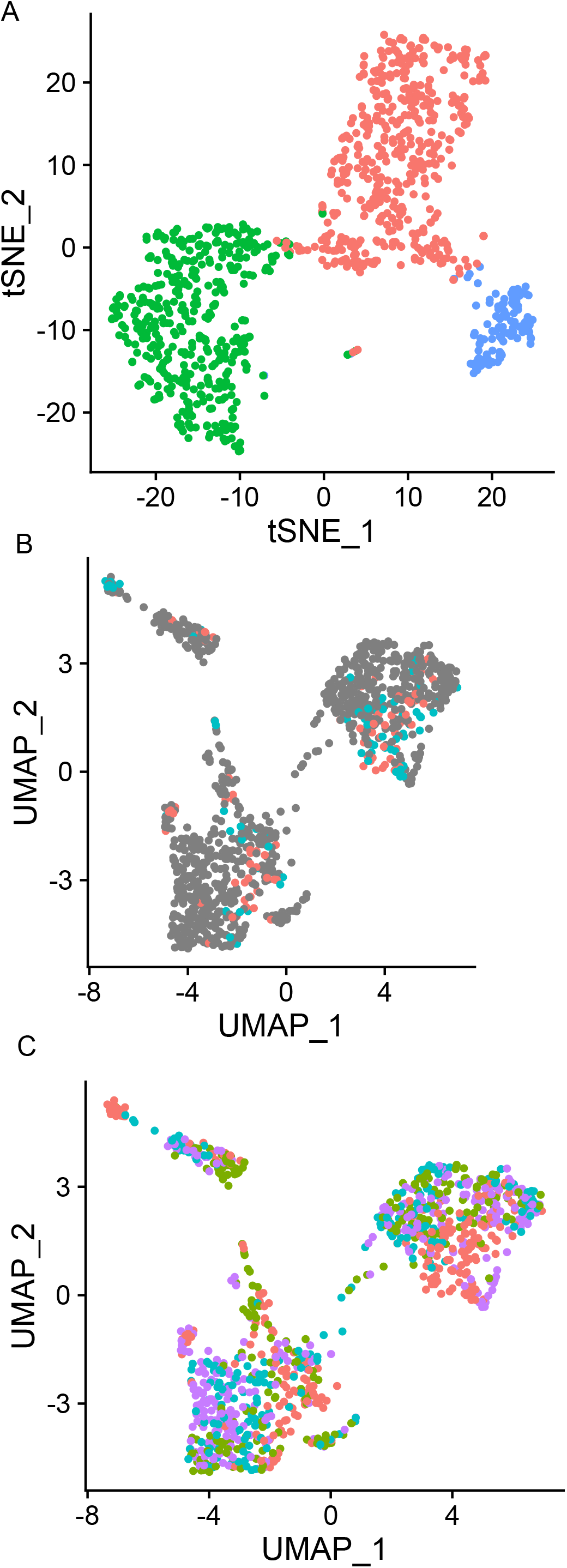
Consistency and reproducibility of scRNA-seq data. A. t-SNE dimensionality reduction of scRNA-seq dataset, illustrating separation of clusters. B. Cells coloured according to location of slice (anterior vs. posterior), depicted in UMAP space. Anterior and posterior cells occupy identical clusters. C. Cells coloured according to animal, depicted in UMAP space, illustrating clustering of cells across animals.

**Figure 1 - figure supplement 2.**
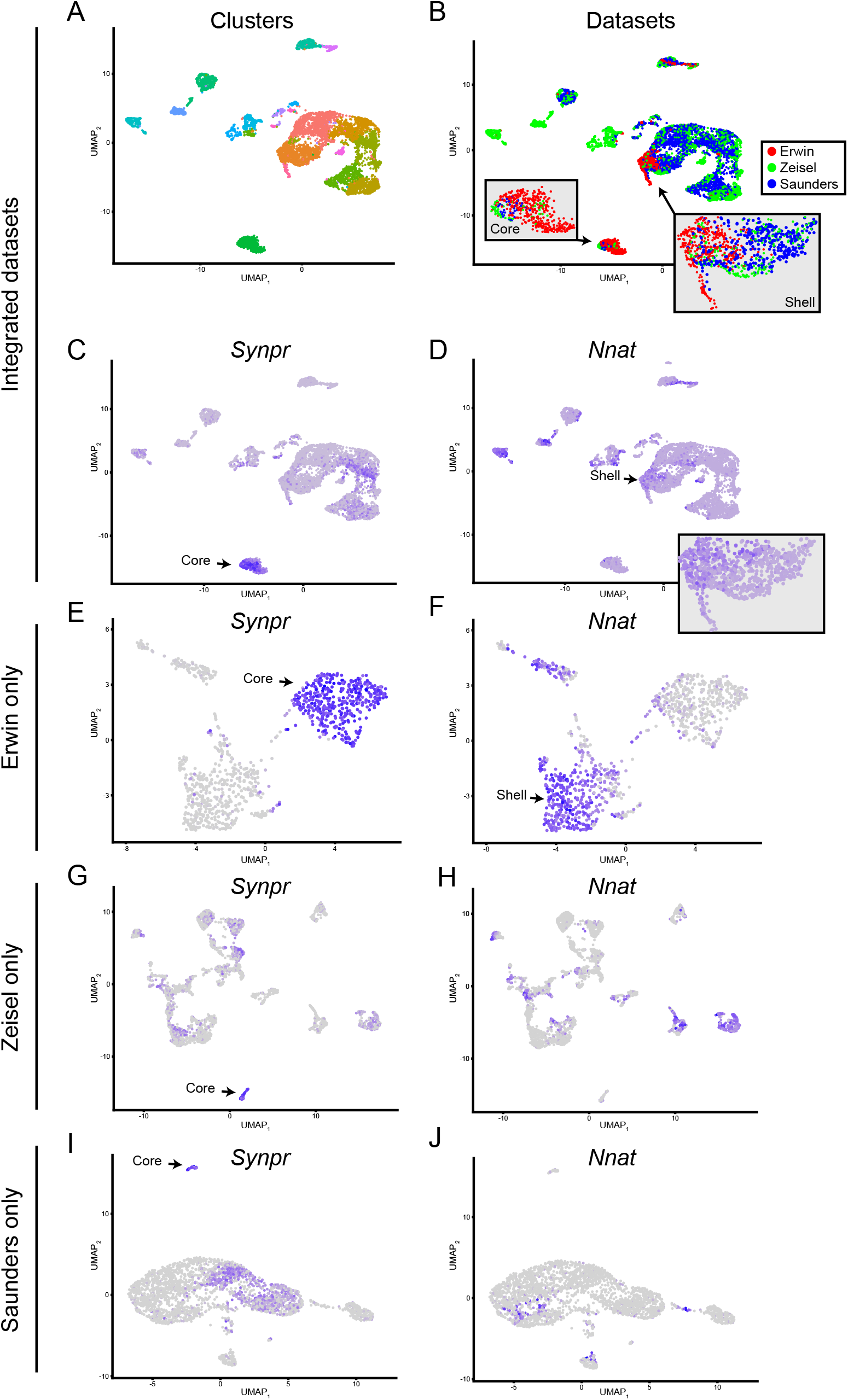
Comparison of new and published scRNA-seq datasets. A. scRNA-seq analysis, integrating the new (“Erwin”) dataset with published data from (Zeisel et al., 2018) and (Saunders et al., 2018) (“Zeisel” and “Saunders”, respectively). Cells in the integrated dataset are depicted in UMAP embedding and coloured by cluster identity. B. As in (A), but coloured according to source dataset. Insets provide expansion of cell types enriched in the Erwin dataset. C. Expression of *Synpr* in the integrated dataset, helping to illustrate the claustral core. D. As in (C), but for *Nnat,* illustrating the claustral shell. E,F. Expression of *Synpr* (E) and *Nnat* (F) in the Erwin dataset only, illustrating their strong enrichment in core and shell claustrum populations, respectively. G,H. As in (E,F), but for the Zeisel dataset. The *Synpr*-expressing (putative claustrum core) cell type is apparent but relatively underrepresented, and the *Nnat*-expressing (putative claustrum shell) cell type is absent. I,J. As in (G,H), but for the Saunders dataset.

**Figure 1 - figure supplement 3.**
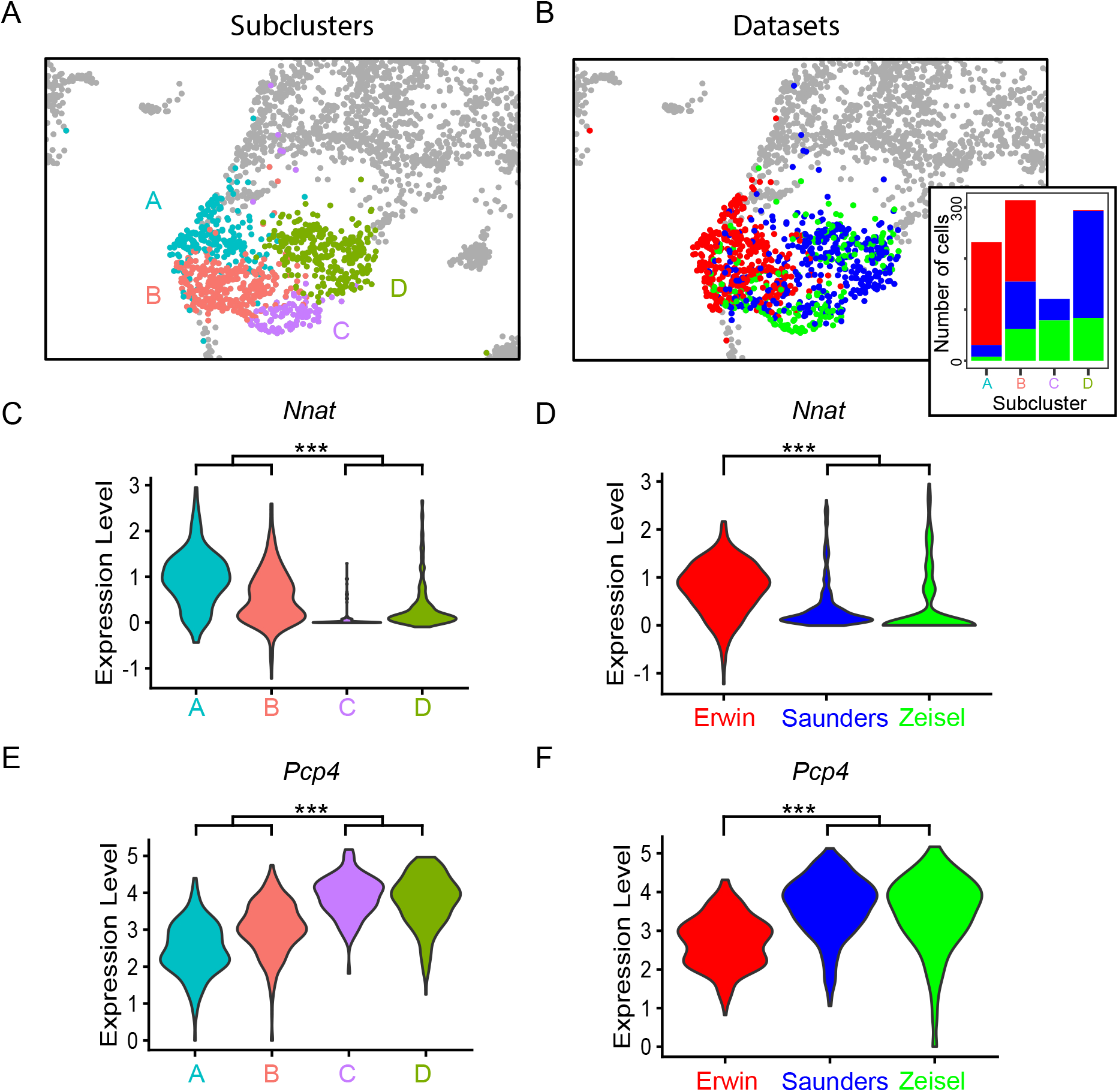
*Nnat* and *Pcp4* differentiate similar cells across datasets. A. Clusters of neurons obtained at a relatively fine clustering resolution, encompassing *Nnat-* expressing claustrum shell neurons from Erwin dataset, as well as similar neurons from published Saunders and Zeisel datasets. B. As in (A), but colored according to dataset of origin. Inset illustrates the number of cells from each dataset on a per-cluster basis. C. Expression of *Nnat* across the clusters in (A). D. As in (C), but expression illustrated by dataset of origin rather than cluster. E,F. As in (C,D), but for *Pcp4*.

**Figure 1 – figure supplement 4.**
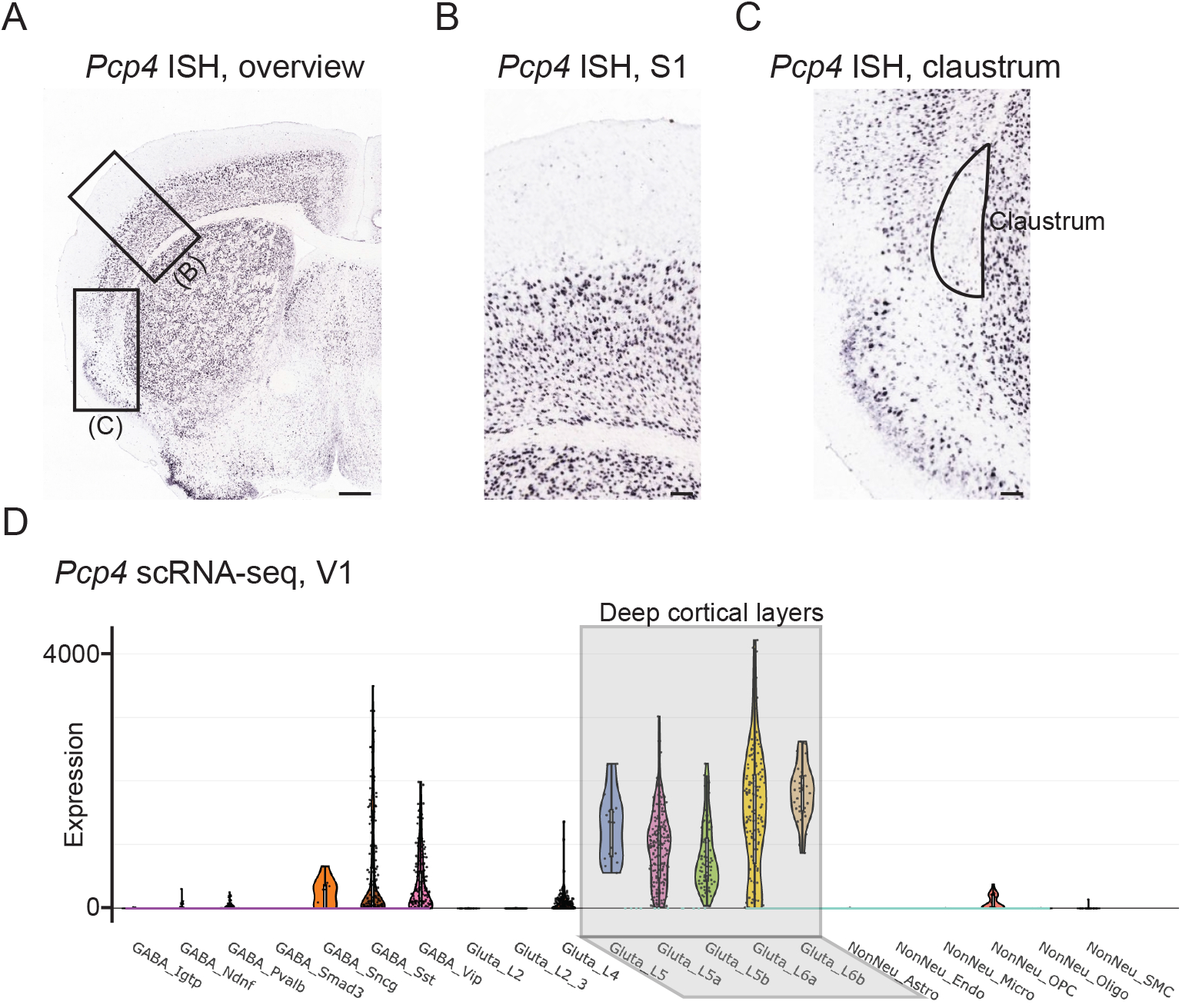
*Pcp4* as a strong and relatively ubiquitous marker of deep cortical layers. A. Chromogenic *in situ* hybridization showing *Pcp4* expression in a coronal section of the mouse brain. Scale bar: 500 μm. B,C. Expansion of the regions highlighted in (A). Note that *Pcp4* exhibits near-ubiquitous expression across the deep cortical layers (e.g., S1), which sharply decreases upon emergence of the claustrum. Data in (A-C) from Allen Mouse Brain Atlas (Lein et al., 2007). Scale bars: 100 μm. D. Single-cell RNA sequencing data from V1 further supporting strong, ubiquitous expression of *Pcp4* across deep cortical layers. Data from (Tasic et al., 2016).

**Figure 2 - figure supplement 1.**
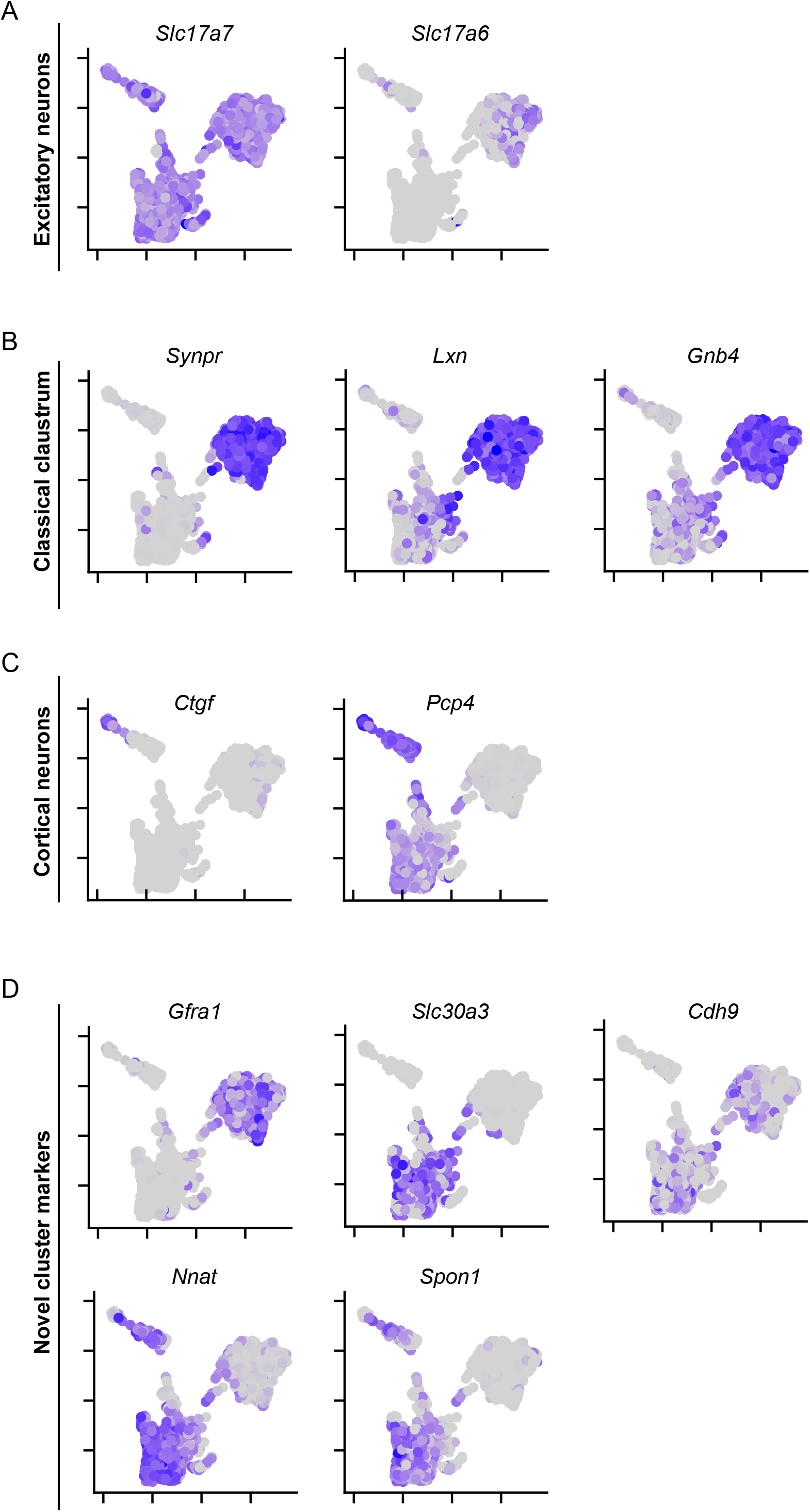
scRNA-seq profiles of mFISH-targeted genes. A. scRNA-seq expression profiles of excitatory neuronal markers *Slc17a7* and *Slc17a6*. Expression depicted in UMAP embedding. B. As in (A), but for previously known claustral marker genes. C. As in (A), but for cortical marker genes. D. As in (A), but for genes identified in this study that were enriched or depleted in a cluster-specific manner.

**Figure 2 - figure supplement 2.**
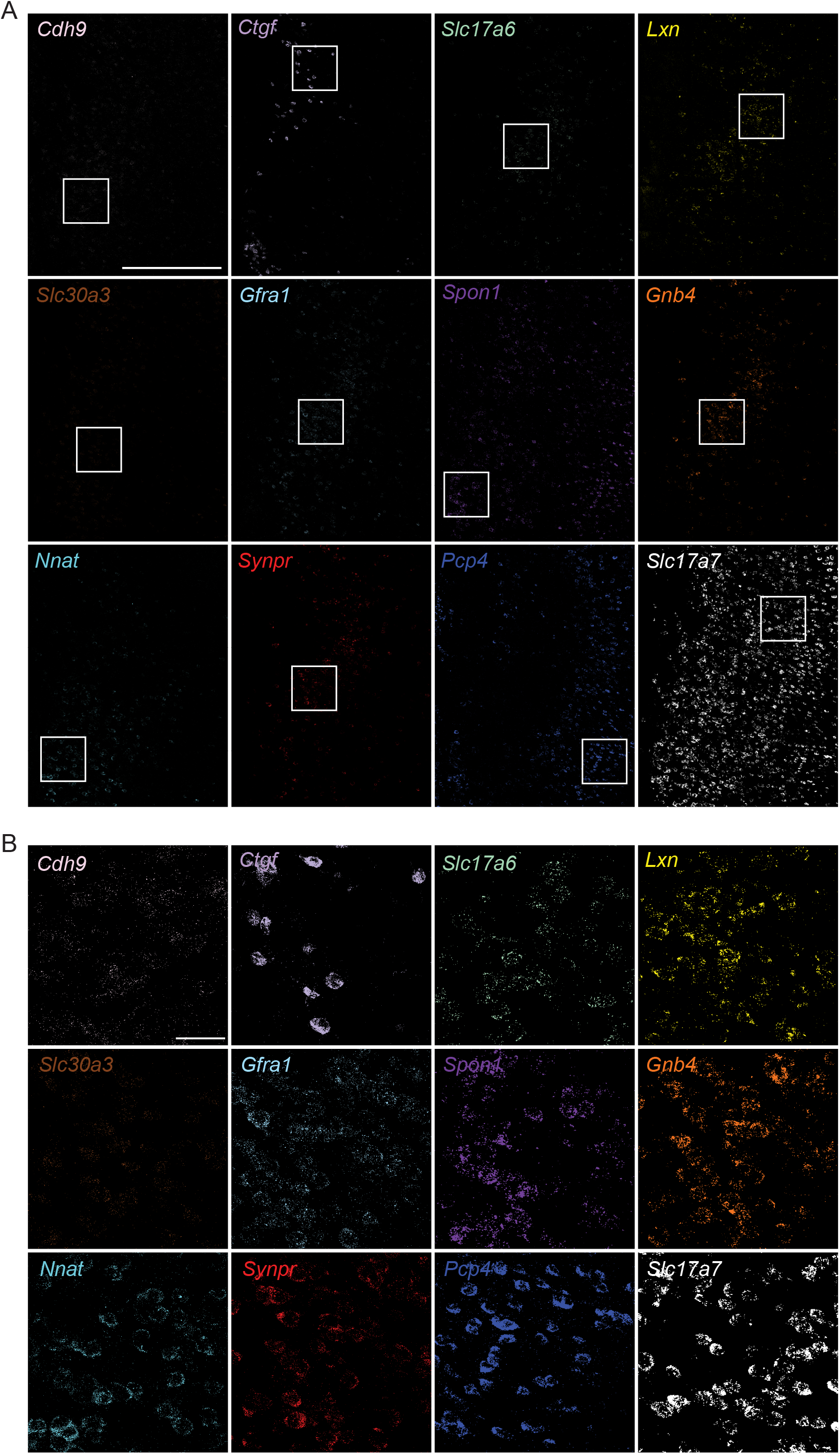
Representative expansions of RNA signals detected via mFISH. A. Overview of signals associated with the 12 genes targeted by mFISH. Scale bar: 500 μm. B. For each gene, expansions of the areas shown in (A). Scale bar: 50 μm.

**Figure 2 – figure supplement 3.**
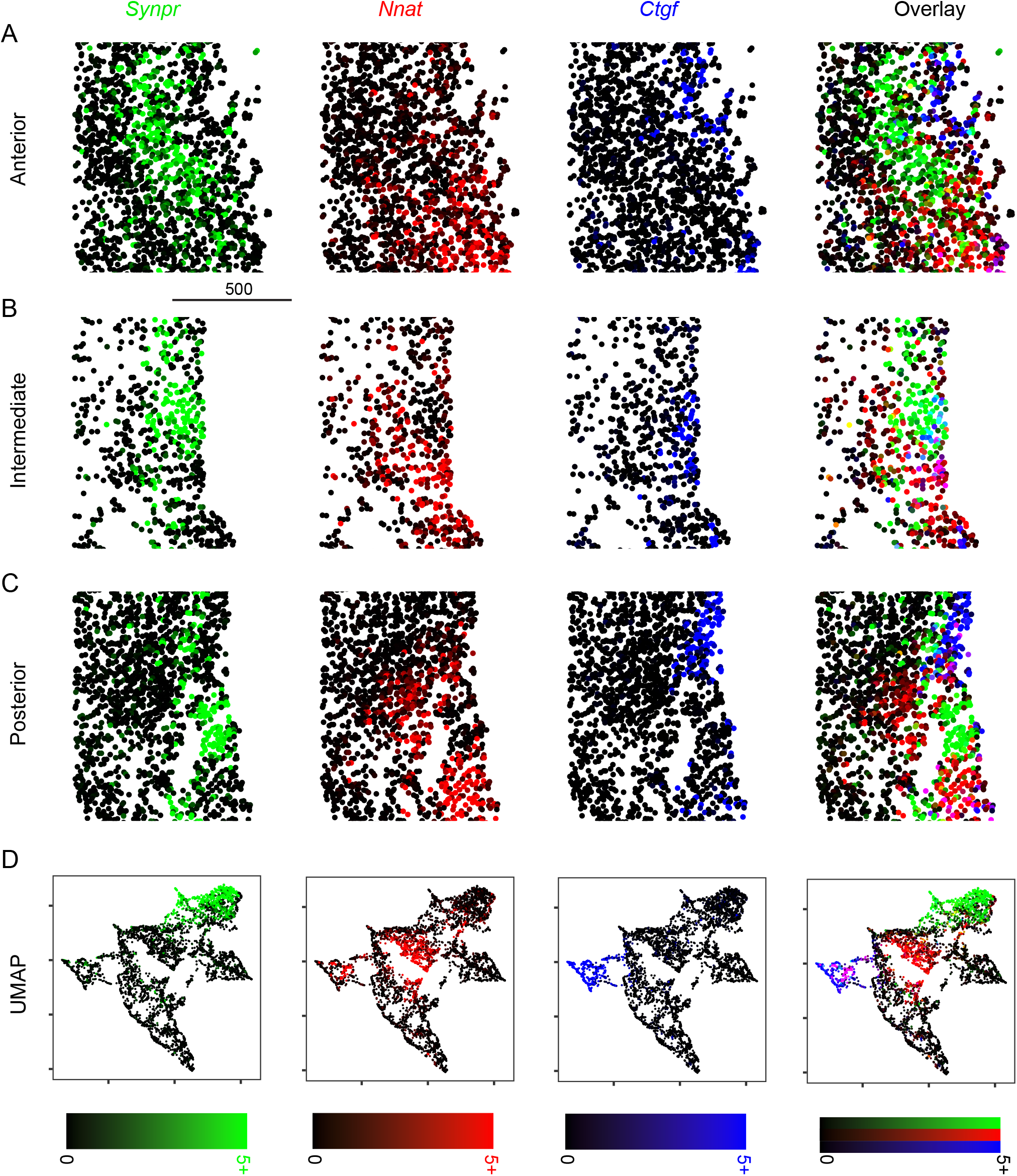
Expression of core, shell, and layer 6 marker genes across the anterior-posterior axis. A. Expression of *Synpr* (green). *Nnat* (red), and *Ctgf* (blue) in anterior sections, as well as three-color overlay. Scale bar: 500 μm. B,C. As in (A), but for intermediate (B) and posterior (C) sections. D. Expression of *Synpr*, *Nnat*, and *Ctgf* in UMAP embedding pooling across all shown sections. Scale bars, illustrating percent area covered (PAC), are provided at bottom for each column.

**Figure 2 - figure supplement 4.**
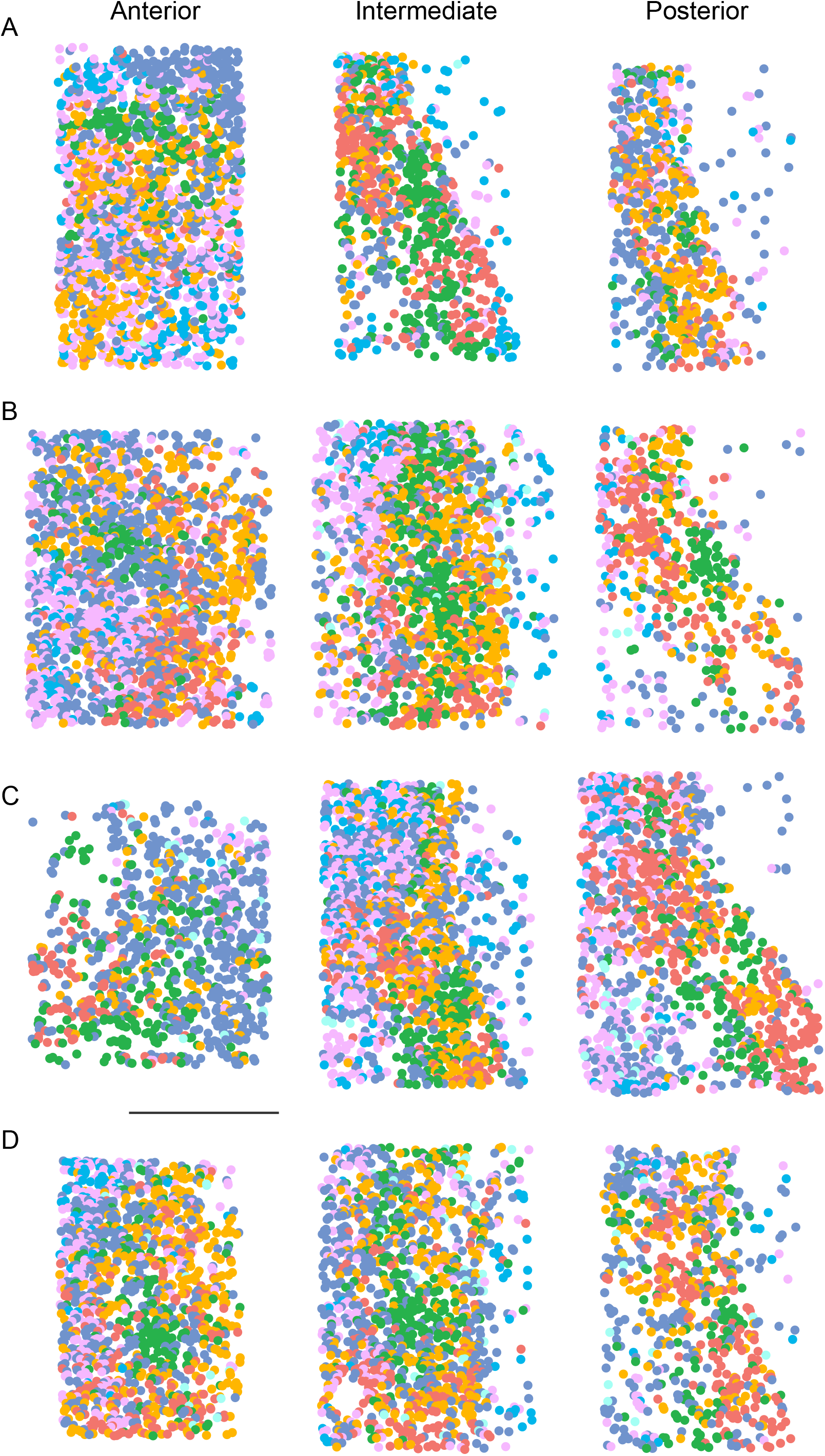
Overview of cellular phenotyping across sections and mice. A. Cellular phenotyping across relatively anterior (left), intermediate (middle), and posterior (right) sections for a replicate animal. B-D. As in (A), but for other replicate animals. Scale bar: 500 μm.

**Figure 2 - figure supplement 5.**
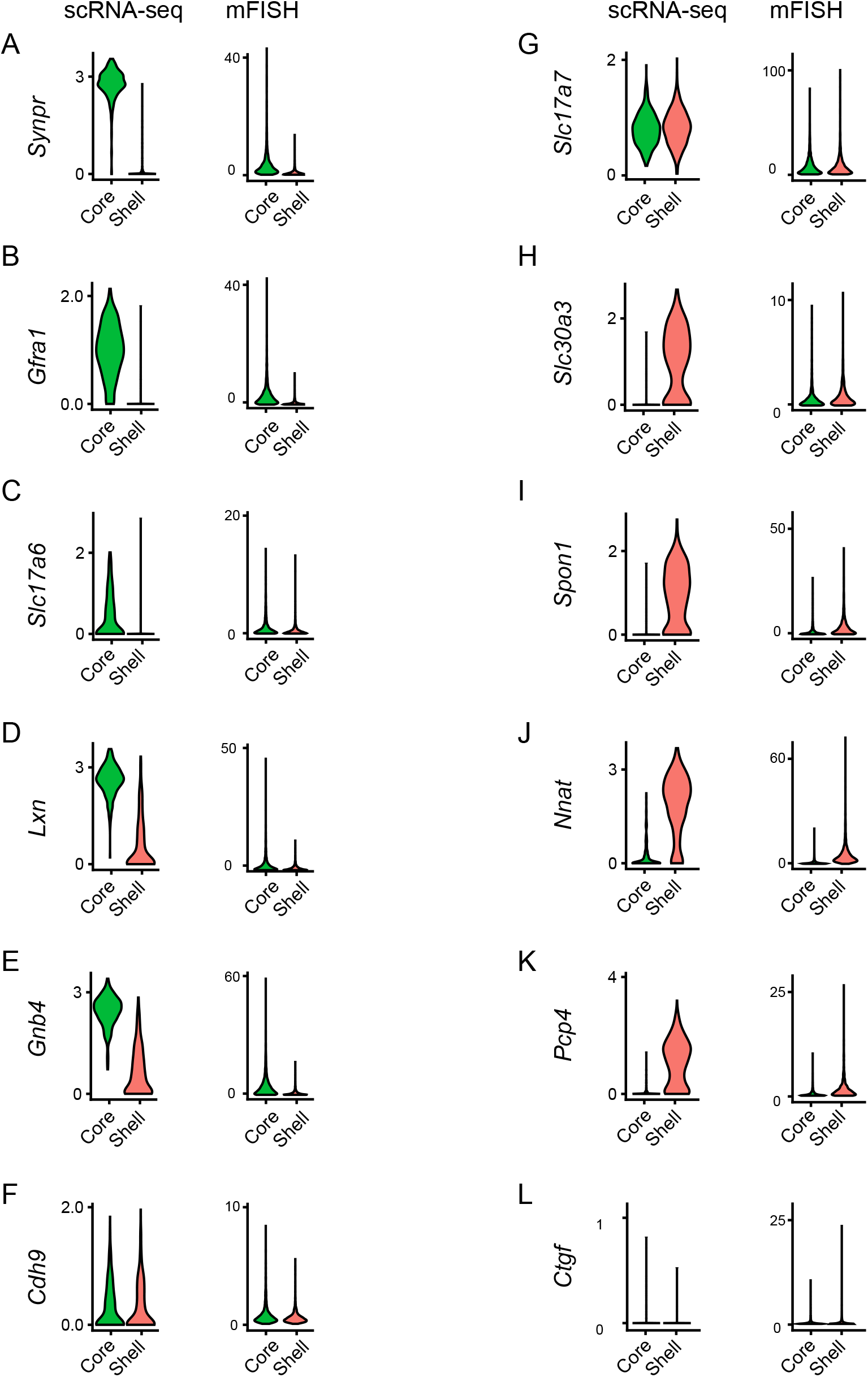
Comparison of gene expression of putative core and shell populations across scRNA-seq and mFISH. A. Violin plots illustrating expression of *Synpr* in scRNA-seq (left) and mFISH (right) in core and shell clusters from both datasets. B-L. As in (A), but for other genes targeted in mFISH.

**Figure 3 – figure supplement 1.**
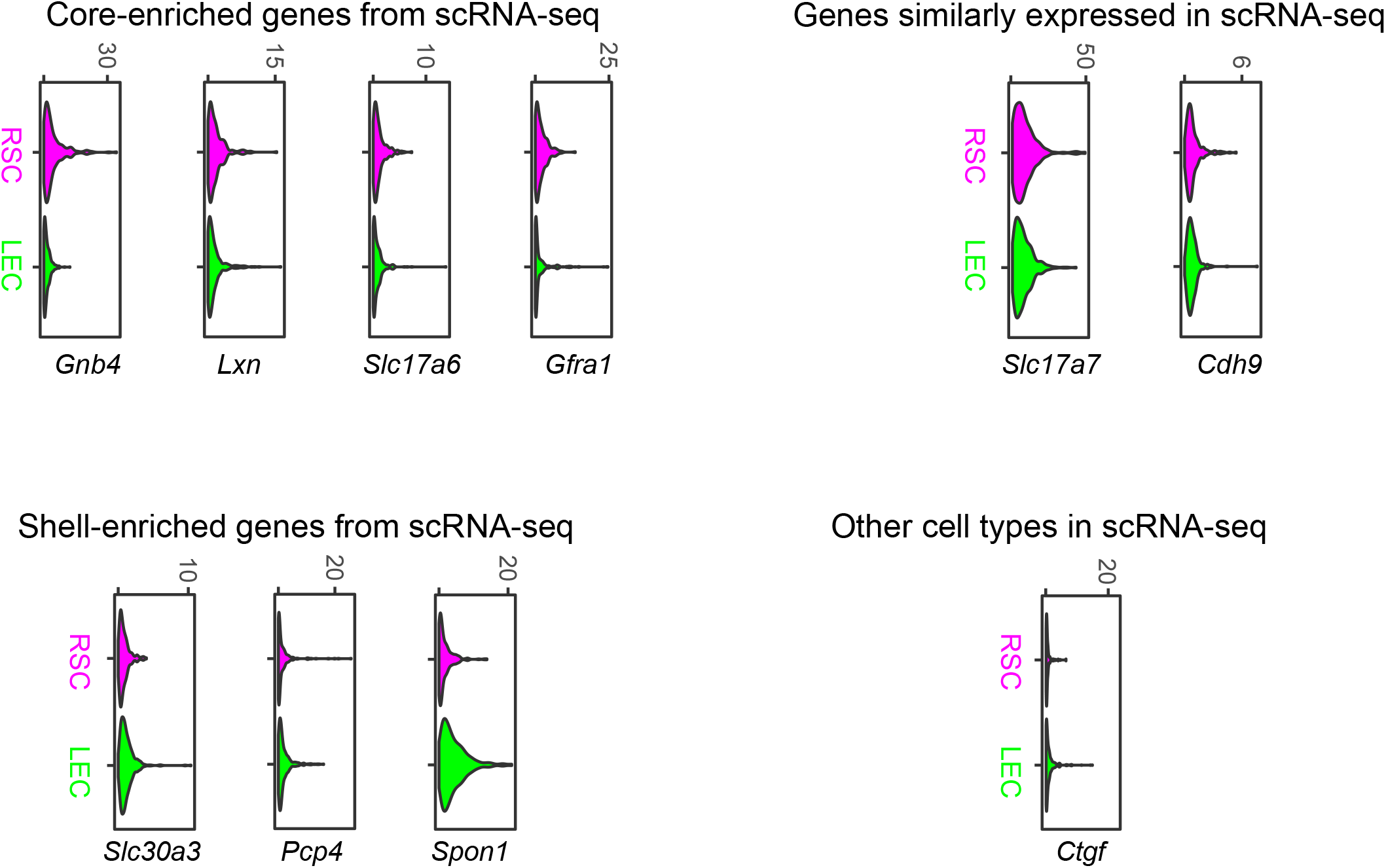
mFISH-derived gene expression profiles for RSC and LEC projecting neurons. Violin plots illustrating expression of genes in mFISH dataset according to projections to either the RSC or LEC. Plots grouped according to scRNA-seq-derived expression profiles.

**Supplementary File 1. List of core-enriched genes and enrichment properties**.

See coreMarkers.txt.

**Supplementary File 2. List of shell-enriched genes and enrichment properties**.

See shellMarkers.txt.

**Supplementary File 3. List of layer 6-enriched genes and enrichment properties**.

See layer6Markers.txt.

**Supplementary File 4. Example mFISH image across fine and coarse spatial scales**.

See mFish.mov.

